# Prophylactic efficacy of riluzole against anxiety- and depressive-like behaviors in two rodent stress models

**DOI:** 10.1101/2020.08.07.242057

**Authors:** Yashika Bansal, Corey Fee, Keith A. Misquitta, Sierra A. Codeluppi, Etienne Sibille, Robert M. Berman, Vladimir Coric, Gerard Sanacora, Mounira Banasr

**Affiliations:** Campbell Family Mental Health Research Institute, Centre for Addiction and Mental Health, Toronto, ON, Canada; Department of Pharmacology and Toxicology, University of Toronto, Toronto, ON, Canada; Department of Psychiatry, University of Toronto, Toronto, ON, Canada; BioHaven Pharmaceuticals, New Haven, CT, USA; Department of Psychiatry, Yale University School of Medicine, New Haven, CT, USA

**Keywords:** Riluzole, Chronic Stress, Prophylactic, Depression, Antidepressants

## Abstract

**Background:** Chronic stress-related illnesses, such as major depressive disorder and post-traumatic stress disorder share symptomatology, including anxiety, anhedonia, and helplessness. Across disorders, neurotoxic dysregulated glutamate (Glu) signaling may underlie symptom emergence. Current first-line antidepressant drugs, which do not directly target Glu signaling, fail to provide adequate benefit for many patients and are associated with high relapse rates. Riluzole modulates glutamatergic neurotransmission by increasing metabolic cycling and modulating signal transduction. Clinical studies exploring riluzole’s efficacy in stress-related disorders have provided varied results. However, the utility of riluzole for treating specific symptom dimensions or as a prophylactic treatment has not been comprehensively assessed.

**Methods:** We investigated whether chronic prophylactic riluzole (~12-15mg/kg/day p.o.) could prevent the emergence of behavioral deficits induced by unpredictable chronic mild stress (UCMS) in mice. We assessed: i) anxiety-like behavior using the elevated-plus maze, open field test, and novelty-suppressed feeding, ii) mixed anxiety/anhedonia-like behavior in the novelty-induced hypophagia test and, iii) anhedonia-like behavior using the sucrose consumption test. Z-scoring summarized changes across tests measuring similar dimensions. In a separate learned helplessness (LH) cohort, we investigated whether chronic prophylactic riluzole treatment could block the development of helplessness-like behavior.

**Results:** UCMS induced an elevation in anhedonia-like behavior, and overall behavioral emotionality that was blocked by prophylactic riluzole. In the LH cohort, prophylactic riluzole blocked the development of helplessness-like behavior.

**Conclusion:** This study supports the utility of riluzole as a prophylactic medication for preventing anhedonia, and helplessness symptoms associated with stress-related disorders.

## Introduction

Chronic stress is the primary risk factor for a variety of psychiatric disorders including major depressive disorder (MDD), bipolar disorder (BPD), schizophrenia, and post-traumatic stress disorder (PTSD) [1–3]. Together, these disorders which share several common symptoms including low mood, anhedonia and helplessness [4] account for one-third of global years of productivity loss due to disability [5]. Notably, the burden of these disorders has substantially increased over the last couple of years due do to the global COVID-19 pandemic [6,7]. Selective serotonin reuptake inhibitors (SSRIs) serve as first-line treatment approaches for several of these disorders [8–10]. Indeed, antidepressant drugs (ADs) have demonstrated efficacy in treating mood and anxiety related symptoms across these disorders and provide neuroprotection in stress-related illnesses [11]. However, they require several rounds of treatment to achieve remission in a majority of patients, thus exhibiting lengthy delays until remission is achieved [12,13]. Moreover, in PTSD only 59% of individuals receiving SSRI treatment responded after fourteen weeks [14]. Other pharmacotherapies for stress-related illnesses include selective noradrenergic reuptake inhibitors, tricyclic antidepressants and anti-adrenergic agents [15–17]. Notably, approximately a third of MDD patients do not achieve remission despite multiple medication trials [18]. The limited therapeutic efficacy of these drugs points to the need for more effective therapeutics or potential strategies to prevent the onset of mental illnesses. Individuals respond differently to stressful life events [3,19–21] with some showing resilience (adapting to the stressful situations), and others showing susceptibility i.e. developing pathological states such as MDD or PTSD. Similar resilience and susceptibility to stress can be found in rodent models of chronic stress exposure [22–25]. Preclinical studies have focused on leveraging these individual differences to address the maladaptive response to stress and the associated symptom emergence by developing pharmaceuticals with prophylactic properties [26–29]. Therefore, several drugs have been investigated for potential secondary prophylactic effects specifically their use in clinical setting to prevent recurrence or relapse. One such drug is fluoxetine but unfortunately, its administration has little to no prophylactic efficacy in rodent stress models [30]. However, pretreatment with serotonin receptor 4 agonists or ketamine showed effective prophylactic properties [31]. Specifically, pretreatment with ketamine, the glutamate (Glu) *N*-methyl-D-aspartate (NMDA) receptor antagonist was shown to attenuate fear response in the contextual fear conditioning [29] and prevent induction of depressive-like behavior in the chronic social-defeat stress, learned helplessness (LH), and chronic corticosterone models [30,32–34]. Although there is data from literature suggesting that glucocorticoid inhibitors [35,36] and beta-adrenergic compounds such as propranolol have antidepressant effect and may be used for prophylactic purposes [37,38] but these data are controversial [35,36,39]. Theoretically, the use of prophylactic drugs could prevent chronic stress-related pathogenesis in individuals and be beneficial in clinical settings, in particular in patients who suffered one depressive or PTSD episode as well as MDD or BPD patients who discontinue their treatment therefore patients that are at high risk for relapse [40–43]. Other populations that could benefit from prophylactic treatment include individuals exposed to acute trauma and who are at high risk for developing PTSD [44]. This is accordance with findings of the literature suggesting that some treatments such as lithium maintenance therapy can increase resilience in clinical MDD and BPD patients [45].

Extensive evidence from both preclinical and clinical studies indicate abnormalities of glutamatergic neurotransmission in the development of psychiatric disorders [32,45–48]. Specifically, alterated regulation of Glu metabolism and neurotransmission and changed expression/function of Glu receptors have been reported in MDD and PTSD patients, as well as in postmortem subjects [49–52]. Similar glutamatergic dysfunctions are found in rodent models of chronic stress [53]. Specifically, stress exposure induce Glu release in corticolimbic brain regions, including the prefrontal cortex (PFC), hippocampus (HPC), and amygdala [54,55]. Elevated or sustained Glu levels induced by chronic stress can causes excitotoxic effects via enhanced extrasynaptic signaling activity [53,56,57]. These alterations have been linked to maladaptive changes in the structure and function of neurons [58,59] and glial cells [60–63]. Glial cells such as astrocytes play a pivotal role in the homeostatic response to the stress-induced increases in glutamatergic neurotransmission [53,62,64,65] as they are responsible for over 80% of Glu reuptake, Glu synthesis and metabolism [66–68]. In light of this, drugs that enhance Glu clearance may prevent the deleterious effects of excess Glu [69–72].

Riluzole, a FDA-approved drug for amyotrophic lateral sclerosis, provides neuroprotection by enhancing synaptic reuptake of elevated excitatory amino acid transporters (EAATs) present on the astrocytes; thereby increasing the ratio of synaptic to extra-synaptic signaling which further promotes neurotrophic factor expression [45,73,74]. Indeed, in chronic stress- or glucocorticoid-exposed rodents, chronic riluzole treatment reversed depressive-like behavior [61,75]. Chronic riluzole administration also attenuated the structural and functional deficits in PFC and HPC as indicated by markers of synaptic Glu signaling and increased levels of brain derived neurotropic factor (BDNF) [61,75]. Riluzole shares neurotrophic effects with conventional antidepressants and its modulation of Glu signaling with rapid-acting antidepressants, such as the NR2B-containing NMDAR antagonist ketamine [76]. Moreover, early open-label studies in MDD patients suggested riluzole possessed antidepressant properties when used either as monotherapy or as adjunctive treatment of monoaminergic antidepressants [77–80]. Further, a randomized, double-blind, placebo-controlled (RDBPC) clinical trial in hospitalised MDD patients showed the combination of riluzole plus citalopram to be significantly more efficacious and to have a faster onset of action compared to placebo plus citalopram [81]. However, the largest randomized placebo-controlled study examining the efficacy of rilzuole augmentation of standard antidepressants in the treatment of MDD did not find evidence of riluzole benefit [77]. Additionally, a recent study of 74 patients diagnosed with PTSD failed to demonstrate efficacy on the Clinicin Administered PTSD Scale, the primary outcome measure in the study [78]. This discrepancy may be due to difference in study design, patient history (i.e. length, severity, prior drug administration) or the timing of treatment intervention in relation to the illness, or the result of under-powered studies. Although these studies clearly show that riluzole is ineffectual in reversing the depressive symptoms in MDD and PTSD, the potential use of riluzole as secondary prophylactic intervention has never been investigated.

Recent studies in rodent models of other pathologies such as spinal cord injury [83] or autoimmune encepthalomyelitis [84] suggest that riluzole treatment might be more effective at reducing behavioral and cellular deficits associated with these models when administered prophylactically. Thus, in this study we investigated if riluzole confers enhanced stress resilience when administered prophylactically, i.e., concurrent with unpredictable chronic mild stress (UCMS), while specifically assessing anxiety-, anhedonia-like behaviors, and overall behavioral emotionality [60,85,86]. In a separate cohort, we also investigated whether chronic preventative riluzole treatment could block the development of helplessness, a core feature of PTSD and other mood disorders. Given that chronic stress and learned helplessness paradigms are well-documented models that appear to recapitulate the behavioral and glutamatergic tripartite synapse deficits akin to those observed in humans with stress-related illnesses [53], we predicted that the Glu-modulating drug riluzole would prevent the emergence of anxiety-, anhedonia-, and helplessness-like behavior. This study is designed to provide preliminary preclinical data supporting the use of riluzole as a secondary prophylactic intervention in stress-related disorder.

## Methods

### Animals

Eight-weeks old male C57BL/6 mice (Jackson Laboratory, Saint Constant, QC) were housed under a 12hr light/dark cycle at constant temperature (20-23 °C) with *ad libitum* access to food and water, except when subjected to light disturbances or food/water deprivation during unpredictable chronic mild stress (UCMS) or learned helplessness (LH) procedures. Following a 2-week facility habituation, animals were single-housed and randomly assigned to 1 of 4 treatment groups: control+vehicle, control+riluzole, UCMS+vehicle, or UCMS+riluzole (*n*=10/group). A separate cohort of 8-week-old male C57BL/6 mice (Jackson Laboratory, Bar Harbor, ME) were habituated to the facility under group-housed conditions throughout the experiment. LH mice were randomly assigned to 1 of 2 treatment groups: LH+vehicle or LH+riluzole (*n*=10/group). All testing was performed during the animal’s light cycle and in accordance with Institutional and Canadian Council on Animal Care (CCAC) guidelines. This study protocol was reviewed and approved by [Animal Care Committee and Centre for Addiction and Mental Health], approval number [ACC725]

### Drug Administration

The riluzole was administered at a dose of ~12-15mg/kg/day p.o in 0.3% saccharin (Sigma Aldrich, St. Louis, MO) in drinking water, known to have minimum toxicity in rodents [87]. The dose was selected based on the previous study aiming to identify the antidepressant-like properties of oral riluzole in the chronic corticosterone mouse model [88]. The average water intake of a mouse weighing 22-30 g is approximately 6-8ml/day [70]. So, based on that the riluzole concentration in drinking water was kept 60ug/ml [88]. Daily drinking was monitored for each animal. Riluzole was dissolved by stirring in the saccharin solution at room temperature. Solutions were changed every 48-72hrs. Mice not treated with riluzole were kept on saccharin alone (vehicle). To assess the prophylactic efficacy of riluzole in UCMS, drug administration commenced on the same day as stressors. To assess preventative efficacy in LH, riluzole commenced 2 weeks prior to the inescapable footshock session. Both cohorts were maintained on riluzole throughout the duration of the experiments, except when fluid deprivation was required for the sucrose and water consumption tests.

### Unpredictable Chronic Mild Stress (UCMS) Procedure

UCMS mice were subjected to 5 weeks of randomized stressors (2-4/day) and maintained for 2 weeks during testing (1-2/day) based on previously validated methods [86,89,90]. Stressors included: forced bath (~2cm of water in rat cage for 15min), wet bedding (30min-1h), aversive predator odor (20min-1h exposure to fox urine), light cycle reversal or disruption, social stress (rotate mice between cages within drug groups), tilted cage (45° tilt up to 2h), reduced space (dividers in cages up to 2h), acute restraint (15-30min in 50mL falcon tube with air/tail holes), bedding change (replacing soiled bedding with clean bedding), bedding or nestlet removal (up to 2h). Control animals underwent regular handling.

### Behavioral Tests

For the UCMS cohort, behavioral testing commenced in week 3. Tests assessed locomotor activity (LA), anxiety-like behaviors in the elevated plus-maze (EPM), open field test (OFT) and novelty-suppressed feeding (NSF) tests, anhedonia-like behavior in the sucrose consumption test (SCT), or both in the novelty-induced hypophagia test (NIH). Physiological deficits were tracked with weekly fur coat assessment during 5 weeks of UCMS exposure and drug treatment. Testing order was randomized, and experimenters were blinded to the treatment history.

#### Locomotor Activity (LA)

On day 21, LA was measured for 20min in a regular home cage (26×15×12cm) using an aerial-mounted video camera. ANY-maze tracking software (Stoelting Co., Wood Dale, IL) extracted distance traveled. LA was assessed to exclude the possibility that changes observed in other behavioral tests were due to bias introduced by treatment-related alterations in ambulatory movement.

#### Elevated Plus Maze

On day 22, animal behavior was assessed in a plus shaped maze consisting of 4 white Plexiglas arms, two open arms (27×5cm) and two enclosed arms (27×5×15cm) situated 55cm high with similar arms faced opposite each other. Animals freely explored the EPM for 10min in a 20-lux room. An aerial-mounted camera/ANY-maze software measured the time spent (sec) and number of entries in open and closed arms.

#### Open Field Test

On day 23, behavior was assessed in an open field (70×70×33cm) placed on the floor. An aerial-mounted camera/ANY-maze software divided the field into 3 concentric square zones of equal area, and measured time spent and number of entries into the innermost zone (40×40cm).

#### Novelty-suppressed Feeding (NSF)

On day 25, following 16h food deprivation, animals were introduced to a novel arena (62×31×48cm) containing a single food pellet. An experimenter blind to treatment conditions measured latency to feed on the pellet during a 12min period. As a control for differences in appetitive drive, latency to feed was then measured in the animal’s home cage during a 6 min period.

#### Novelty-Induced Hypophagia (NIH)

On days 25-26, animals were habituated to a daily liquid reward (1mL of 1/3 v/v sweetened condensed milk). On day 26, latency to drink the reward was measured in the home cage as a control for potential treatment effects on appetite or activity. The following day, the animals were moved to a new cage (26×15×12cm) in a brightly lit (500-600 lux) room and the latency to drink the milk was measured over a 6min trial as a measure for hyponeophagia.

#### Sucrose Consumption Test (SCT)

On days 27-28, home cage water bottles were switched to sucrose solution (1%; Sigma, St. Louis, MO) ± riluzole. Animals were habituated to sucrose for 48h and then fluid deprived overnight (~14h). On day 29, sucrose intake (mL) was measured over a 2h test. After the SCT, animals were returned to regular solutions (saccharine±riluzole) for 24h. The same test was repeated the next day i.e. fluid deprivation (14h) and water intake (1h) to control for potential treatment effects on fluid intake.

#### Fur Coat Assessment

The coat state of each animal was assessed on 7 anatomical areas by attributing a score of 0, 0.5, or 1 for each, from maintained to unkempt. Coat assessment was scored by an experimenter blind to treatment conditions every week for 5 weeks of UMCS.

#### Z-score calculations

The z-score were calculated as in [87]. Briefly the z-score of each test was determined using average and standard deviation of the control group (no stress + vehicle group). Z-anxiety score was calculated by averaging the z-scores of open field, elevated plus maze and NSF while the z-anhedonia score was calculated by averaging the z-scores of NIH and sucrose consumption tests. Z-emotionality scores were calculated by averaging all of the tests

### Learned Helplessness (LH) Procedure

A separate cohort of animals was habituated to a shuttle box (Med Associates, St. Albans, VT) for 5min and underwent a session consisting of 60 randomized, unpredictable, and inescapable footshocks (0.35 mA) administered over ~1h. Two days after the footshock session, mice were tested in the active avoidance (AA) paradigm. Mice underwent 30 trials of randomized footshocks (0.35 mA). Each trial lasted a maximum of 1min and included a minimum of 5sec before each footshock. Then, the door separating 2-halves of the chamber was opened (2sec before footshock initiation) and remained open for the duration of the shock. The first 5 trials required one chamber crossing (FR1) to terminate the footshock, whereas the next 25 trials required two crossings (FR2). The AA session lasted ~30min, during which the number of escape failures was recorded.

### Statistical analysis

All data was analyzed using the Statistical Product and Service Solutions software (SPSS; IBM, North Castle, NY). Based on previously validated methods [91,92], parameters were z-scored and averaged across tests reflecting anxiety-(EPM, OFT, NSF, NIH) and anhedonia-like behavior (NIH, SCT), as well as overall emotionality (all previous + fur coat state in the last week). Normalized z-scores integrate behavioral tests measuring similar phenotypes to increase the consistency and reliability of these characteristically inconsistent tests [89,91,92]. Behavioral data was analyzed using a two-way (stress x drug) analysis of variance (ANOVA), followed by Bonferroni-corrected *post-hoc* comparisons. Weekly coat state measures were analyzed using repeated-measures ANOVA. AA escape failures were analyzed using *student’s t-test* comparing data between vehicle and riluzole groups. Data is presented as mean ± standard error of the mean (SEM).

## Results

### Riluzole prevents the emergence of UCMS-induced anhedonia-like behaviors

We first sought to determine whether chronic prophylactic riluzole treatment (i.e., administered concurrent with UCMS) could block the emergence of behavioral deficits induced by UCMS. Mice were subjected to 3 weeks of daily handling (control) or UCMS ± vehicle or riluzole and tested for 2 weeks, including for LA and in the OFT, EPM, NSF, NIH, and SCT (**Fig 1*a***). Fur coat deterioration was tracked weekly (**Fig. 1*a***). In the LA assessment, distance travelled in a home cage-like environment was not significantly affected by stress (*F_1,36_*=0.025; *p*=0.87), drug treatment (*F_1,36_*=0.183; *p*=0.67), or stress*drug interaction (*F_1,36_*=3.02; *p*=0.09), confirming that subsequent tests were not biased by alterations in ambulatory movement (*not shown*).

**Fig. 1.**
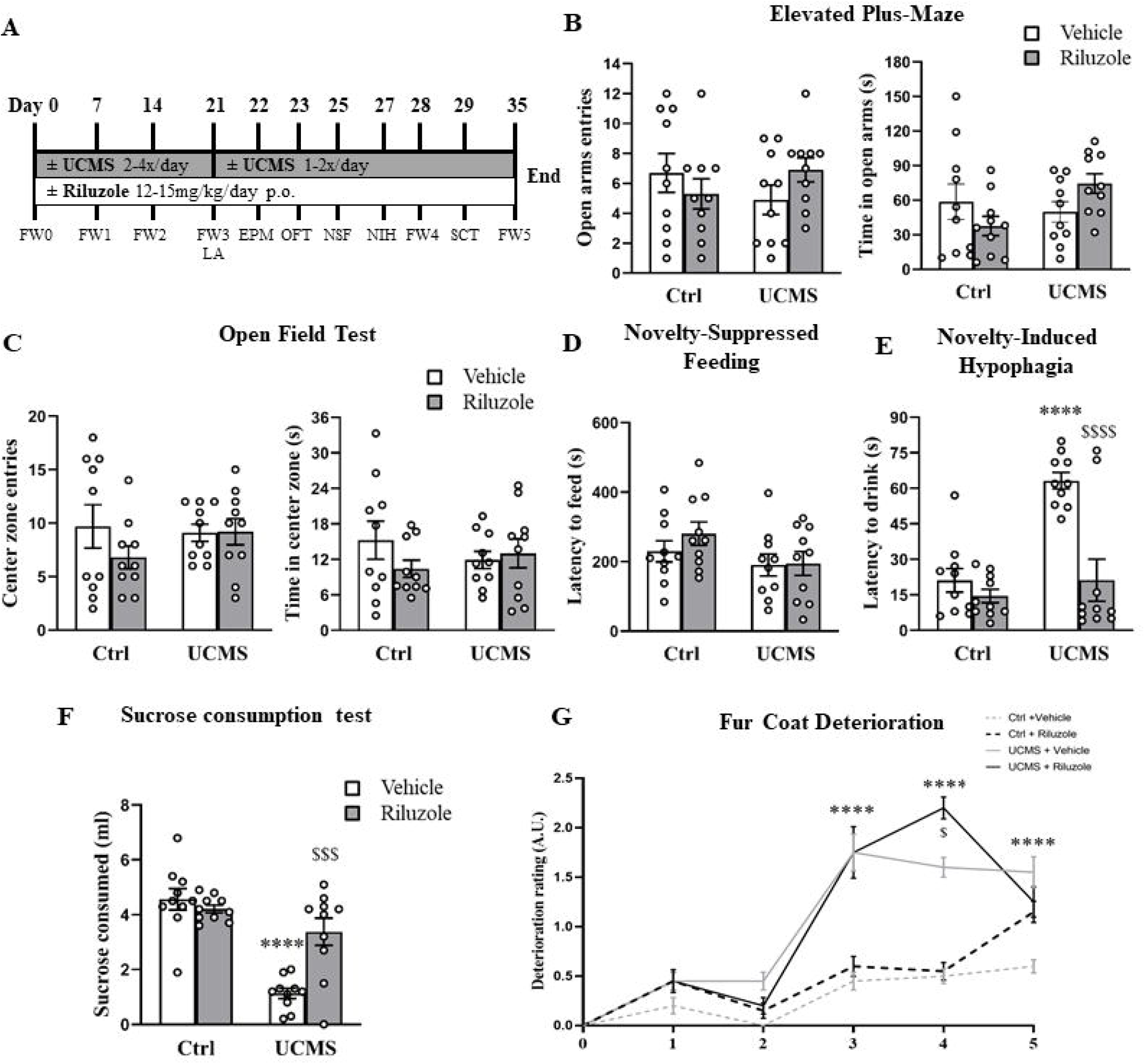
Riluzole prevents the emergence of anhedonia-like behaviors induced by unpredictable chronic mild stress (UCMS).

(**A**) study design for behavioural analysis of Fur/Weight (FW0-5), Locomotor Activity (LA), Elevated-Plus Maze (EPM), Open Field Test (OFT), Novelty-Suppressed Feeding (NSF), Novelty-Induced Hypophagia (NIH), and Sucrose Consumption Test (SCT) beginning on day 21 of Control (Ctrl)/UCMS ± Vehicle (Veh)/Riluzole 13.2mg/kg/day p.o. (*n* = 10/group). (**B**) Number of open arm entries (left) and time spent in open arms in the EPM. (**C**) Number of center zone entries and time spent in the center zone in the OFT. (**D**) Novel environment latency to feed in the NSF. (**E**) Novel environment latency to drink a milk reward in the NIH. (**F**) Sucrose consumed in the SCT. (**G**) Weekly fur coat deterioration ratings. *\*\*\*\*p*<0.0001 UCMS + Veh vs. Ctrl + Veh; *^$$$$^p*<0.0001, *^$$$^p*<0.001, *^$^p<0.05* UCMS + Riluzole vs. UCMS + Veh.

In the EPM, a two-way ANOVA of open arm entries revealed no main effect of stress (*F_1,36_*=0.009; *p*=0.92), drug (*F_1,36_*=0.084; *p*=0.77), or a stress*drug interaction (*F_1,36_*=2.68; *p*=0.11; shown in **Fig.1.*b***). For time spent in the open arms, a two-way ANOVA revealed a significant stress*drug interaction (*F_1,36_=4.49; p*=0.041; shown in **Fig. 1.*b***); however, group-wise differences did not survive *post-hoc* correction.

In the OFT, no significant differences were found for the effects of stress, drug treatment, or their interaction on the number of entries or time spent in the center zone (shown in **Fig. 1.*c***).

In the NSF, a two-way ANOVA revealed a trend towards a main effect of stress on latency to feed in the novel environment (*F_1,36_*=3.71; *p*=0.061; shown in **Fig. *1.d***). However, this may have been confounded by an UCMS-induced increase in appetitive drive, since feeding latency in the home cage control test was significantly reduced for both stress groups (*F_1,36_*=12.68; *p*=0.0011) (CTR+vehicle: 71.7±10.9; CTR+riluzole: 77.1±8.3; UCMS+vehicle: 49.3±6; UCMS+riluzole: 42.4±5.5). The UCMS paradigm includes brief food deprivation experiences which may bias animals towards increased motivation for feeding after longer deprivation periods such as with the NSF. Indeed, in the NIH, animals were not food-deprived and underwent a similar assessment for reward approach. In the NIH, two-way ANOVA detected a significant main effect of stress (*F_1,36_*=19.29; *p*<0.001), drug treatment (*F_1,36_*=19.13; *p*<0.001), and a stress*drug interaction (*F_1,36_*=10.15; *p*=0.003) for latency to drink the milk in a novel environment (shown in **Fig. *1.e***). *Post-hoc* analysis revealed that this was driven by 3-fold increase in latency to drink induced by UCMS (*p*<0.0001) that was prevented by riluzole (*p*<0.0001). Home cage latency was not affected by stress, drug treatment, or their interaction, suggesting that novel environment differences in approach latency were not due to appetite (CTR+vehicle: 23.2±4.8; CTR+riluzole: 29.2±5.2; UCMS+vehicle: 27.4±11.2; UCMS+riluzole: 46.1±9.4).

In the SCT, two-way ANOVA revealed a significant main effect of stress (*F_1,36_*=39.61; *p*<0.0001), drug treatment (*F_1,36_*=7.880; *p*=0.008), and stress*drug interaction (*F_1,36_*=14.76; *p*=0.0005) for 1hr sucrose consumption following overnight deprivation (shown in **Fig. 1*f***). Further *post-hoc* analysis revealed a significant decrease in sucrose consumption (*p*<0.0001) that was prevented by riluzole (*p*=0.0002). After a 24hr recovery, mice were again deprived and measured for water consumption under identical conditions. Although we detected a main effect of stress on water consumption (*F_1,36_*=5.89; *p*=0.02), reflecting reduced water intake among UCMS-exposed animals, group-wise differences did not survive *post-hoc* analyses, confirming that sucrose consumption differences were reliably due to stress and/or drug treatment effects (CTR+vehicle: 1.6±0.2; CTR+riluzole: 1.5±0.1; UCMS+vehicle: 1.1±0.1; UCMS+riluzole: 0.9±0.2).

Repeated-measures ANOVA of weekly fur coat state assessments from week 0 to 5 revealed significant main effect of stress (F_1,36_= 125.46; p<0.0001), time (F_5,180_= 111.3; p<0.0001), stress*time interaction (F_5,180_= 32.03; p<0.0001) and stress*drug*time interaction (F_5,180_= 5.023; p=0.002), but no main effect of drug and no drug*stress or drug*time interactions. *Post-hoc* analysis showed that UCMS induced persistent fur coat deterioration starting on week 3 which was also significant for each following week (p<0.0001; Fig 1g) and not modified by riluzole treatment.

### Subsequent z-score normalization confirms that riluzole prevents the emergence of anhedonia-like behavior, and composite emotionality-like behavior in UCMS-exposed mice

One issue with rodent chronic stress paradigms and classical testing methods, noted by us and others, is insufficient reliability to consistently measure depressive-like features across tests, cohorts, and between labs [90,93,94]. To address this, we employed multiple tests assessing common dimensional phenotypes (anxiety- and anhedonia-like behavior, and emotionality) and analyzed both individual test parameters and normalized z-scores averaged across tests, based on validated methods [91].

Two-way ANOVA of normalized Z-anxiety scores (reflecting EPM, OFT, NSF, NIH) revealed a marginally significant stress*drug interaction (*F_1,36_*=3.996; *p*=0.0532; shown in **Fig. 2.*a***) with no main effect of stress or drug. However, post-hoc analysis did not reveal group differences. Z-anhedonia (reflecting NIH, SCT) was significantly affected by stress (*F_1,36_*=36.74; *p*<0.001), drug treatment (*F_1,36_*=5.899; *p=*0.0203), and their interaction (*F_1,36_*=9.032; *p*=0048), wherein UCMS induced an elevation in anhedonia-like behavior (*p*<0.0001) that was prevented by riluzole (*p*=0.0029; shown in **Fig. 2.*b***). Summary of behavioral emotionality via Z-emotionality scores analysis (Z-anxiety and Z-anhedonia) revealed significant effects of stress (*F_1,36_*=11.93; *p*=0.0014), drug (*F_1,36_*=2.68; *p*=0.0109) and a stress*drug interaction (*F_1,36_*=11.46; *p*=0.0017; shown in **Fig. 2.*c***). Overall, behavioral emotionality was significantly elevated by UCMS (*p*=0.0001) and prevented by riluzole treatment (*p*=0.0065).

**Fig. 2.**
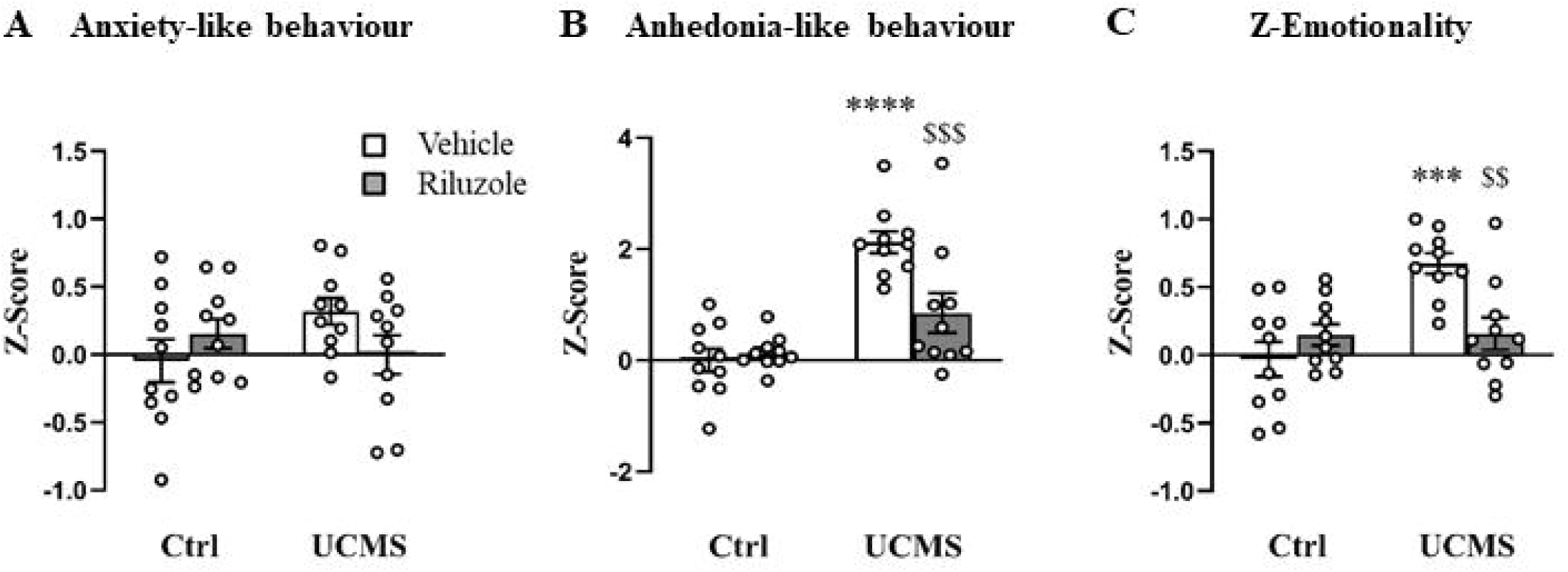
Z-score normalization reveals that riluzole prevents the emergence of anxiety- and anhedonia-like behaviors, and emotionality-like behavior induced by unpredictable chronic mild stress (UCMS). (**A**) Z-anxiety scores (reflecting elevated-plus maze, open field test, novelty-suppressed feeding, and novelty-induced hypophagia behavioral performances). (**B**) Z-anhedonia scores (reflecting novelty-induced hypophagia, sucrose consumption test performances). (**C**) Z-emotionality (reflecting all previous plus fur coat deterioration). (*n*=10/group) \*\*\*\**p*<0.0001, \*\*\**p*<0.001 UCMS + Veh vs. Ctrl+Veh; *^$$$^p*<.001, *^$$^p*<.01 UCMS + Riluzole vs. UCMS + Veh.

### Riluzole prevents the development of learned helplessness

Next, we investigated whether preventative riluzole treatment could block the development of helplessness-like behavior induced by inescapable footshock. A separate cohort of mice (*n*=10/group) were administered riluzole or vehicle for 2 weeks and submitted to unpredictable inescapable footshocks as part of the LH protocol (shown in **Fig. 3.*a***). In the AA paradigm, *t-test* analysis revealed a significant effect of drug treatment on the number of escape failures (*t*=4.55, df=18; *p*=0.0001), wherein compared to vehicle, riluzole decreased the number of escape failures (shown in **Fig. 3.*b***).

**Fig. 3.**
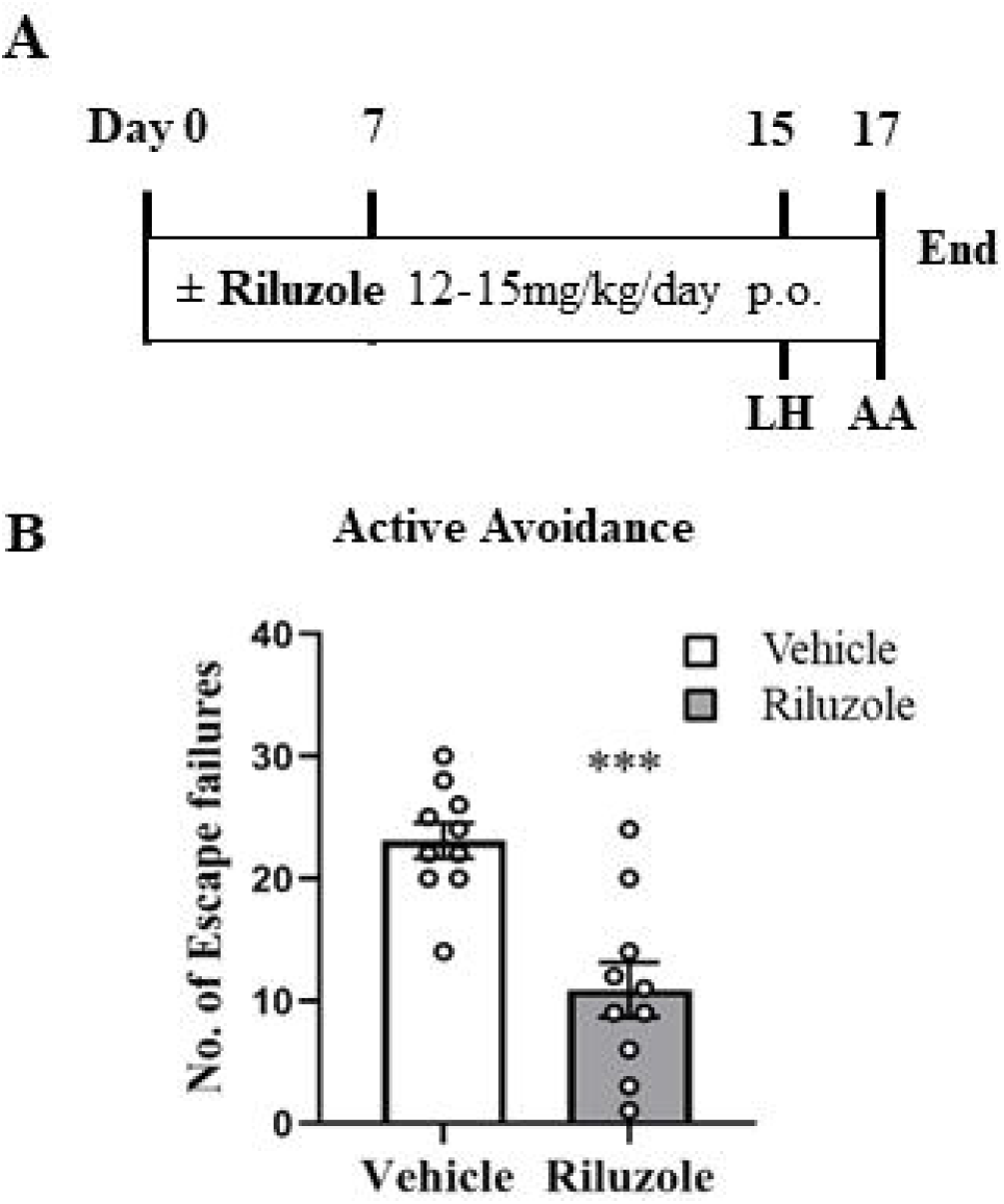
Riluzole blocks the development of learned helplessness (LH). (**A**) Study design for chronic preventative riluzole treatment prior to learned helplessness conditioning including 60 inescapable footshocks with active avoidance (AA) testing 2 days later. (**B**) Number of escape failures in the active avoidance session (*n*=10/group). \*\*\**p*<.001 Riluzole vs. Vehicle (Veh).

## Discussion

This study provides evidence that chronic prophylactic riluzole treatment can increase stress resilience and prevent the development of specific behavioral deficits induced by UCMS and LH. UCMS and LH are relevant rodent models of stress which induce behavioral deficits, such as anxiety, anhedonia and helplessness, commonly associated with stress-related illnesses including depression and PTSD [95–97]. Animals exposed to UCMS exhibited elevated anhedonia-like behaviors in the novelty-induced hypophagia and sucrose consumption tests, which were prevented by chronic concomitant riluzole treatment. In this study, UCMS didn’t induce anxiety-like behaviors making it difficult to assess the effects of riluzole treatment in these tests. Overall, z-score normalization confirms that riluzole prevents UCMS-induced elevation of behavioral emotionality, without affecting the physical state of the animals as we found no blockade of the UCMS effects on coat state in riluzole treated-animals. In a separate cohort, we also demonstrated that 2 week treatment with riluzole prior to inescapable footshocks blocked the development of helplessness-like behavior in the AA test. These findings suggest that treatment with the Glu-modulating drug riluzole may enhance stress resilience and prevent the development of anhedonia and helplessness.

Chronic stress models such as UCMS or chronic restraint stress are classically used to model anxiety and anhedonia-like behavior in rodents. However some tests such as open field or elevated plus maze have been proven to be inconsistent for detecting the anxiogenic effects of UCMS [90,98]. Similar variations have been reported in our previous lab papers [86,89,90,99] and outside labs in mice, rats and different mouse strains as well [98,100–105]. However, topdown dimension reduction via Z-scoring consistently increases the sensitivity and reliability of behavioral tests [91]. Although this approach may improve the quality of conclusions about preclinical behavior changes, it is also important to mention that repeated testing is itself a source of variability [106]. In this study, EPM, OFT, or NSF tests failed to individually detect significant elevation of anxiety-like behavior and also in across tests using the z-anxiety score. In contrast, UCMS induced robust anhedonia-like behaviors in both the NIH and SC tests, confirming previous findings from our lab [60,85,107] and others [108]. In those studies the behavioural effect of chronic stress on anhedonia appear after several weeks of chronic stress exposure (2-3 weeks minimum). This effect was prevented by treatment with riluzole in individual tests and across multiple tests (z-anhedonia score). UCMS also induced coat state deterioration that was unaffected by riluzole. This is consistent with the evidence that fur coat quality tracks the induction of stress-related deficits, but does not respond to most antidepressant treatments [86,90]. Finally, in consideration of the potential sedative effects of high doses of riluzole [74], we confirmed that behavioral outcomes were not influenced by locomotor changes, in agreement with previous studies using similar dosing regimens [61,88]. Altogether, our results demonstrate that chronic riluzole treatment prevents the development of UCMS-induced anxiety and anhedonia deficits.

The LH paradigm models helplessness-like behavior associated with exposure to inescapable and uncontrollable stressors [109], and are used to assess the predictive validity of antidepressant treatment [110]. The acute/intense LH stressor may be more closely related to PTSD given that it reproduces aspects of avoidance, anxiety, and hyperarousal [111]. In rodents, escape failures in the active avoidance paradigm reflect helplessness-like behavior and was shown to partially correlate with other dimensions including anhedonia-like behavior [112]. In a previous study, we demonstrated that chronic riluzole administration reversed anhedonia- and helplessness-like behavior in UCMS-exposed rats [61]. The antidepressant action of riluzole was also shown in other models, such as following olfactory bulbectomy [113] or chronic corticosterone treatment [88]. Here, we identified for the first time that chronic pre-treatment with riluzole significantly reduced the number of escape failures in the active avoidance test. This is in accordance with another study demonstrating that pre-treatment with anti-glutamatergic agents with antidepressant properties such as ketamine, an antagonist of the NMDA receptor prevented behavioral deficits induced by social defeat and LH [88]. Our results suggest that prophylactic administration of riluzole may prevent the development of helplessness-like behavior.

Although outside the scope of this study, we can speculate on the potential cellular mechanisms underlying the prophylactic effects of chronic riluzole administration in preventing the effects of stress. Riluzole’s neuroprotective and antidepressant actions have been associated with several neuronal and non-neuronal cellular mechanisms including modulation of glutamate release and enhancement of glutamate uptake and neurotrophic factors [61,74,88]. In previous work, we demonstrated that riluzole exerts its antidepressant-like efficacy through increased glutamate transporter-1 (GLT-1) expression as well as increased glutamate cycling in neurons and glial cells in chronically stressed and non-stressed rats [61,114]. Anti-glutamatergic effects of riluzole were also shown to involve inhibition of the release of glutamate through a blockade of voltage-activated sodium channels [115,116]. Finally, neuroprotective effects of riluzole have also been associated with increased BDNF expression *in vivo* [73,88,117,118] and *in vitro* by neurons and astrocytes [73,119]. Interestingly, prophylactic effects of riluzole were also demonstrated in aging studies, where pretreatment was reported to prevent cognitive decline in aged rodents through increased GLT-1 expression and maintenance neuronal morphology/complexity [120,121]. More recently, prodromal riluzole treatment was described to induce long-term pro-cognitive effects in a genetic mouse model of Alzheimer’s disease. This effect was associated with attenuated glutamatergic neurotransmission which was observed even six months after riluzole treatment discontinuation [122]. Collectively, these findings implicate Glu homeostasis, modulation, and subsequent neuroprotective or neurotrophic effects in the prophylactic and therapeutic actions of riluzole.

Initial open-label clinical studies have observed significant symptom improvement following oral riluzole administration in patients with treatment resistant depression [77,79]. Follow-up clinical studies have confirmed or found limited efficacy of riluzole monotherapy/ adjunctive therapy to monoaminergic antidepressant or ketamine [74,82,123–125]. While some of these trials presented promising evidence for riluzole antidepressant action, double-blind placebo studies have demonstrated that adjunctive riluzole in MDD and riluzole monotherapy in bipolar depression, failed to induce clinically relevant improvement of depressive symptoms [124,125]. It is possible that the reported mixed efficacy of riluzole in clinical studies is due to variance in treatment protocol or patient characteristics. For example, extension of an 8-week open-label riluzole trial to 12 weeks achieved response in one-fourth of non-responders [126]. Riluzole may also be more useful in patients with deficits in specific biomarkers. Indeed, it was demonstrated that patients with lower BDNF plasma concentrations at baseline (pre-treatment) show greater treatment response to riluzole [127]. Altogether, these clinical studies suggest that riluzole treatment is ineffectual in reversing depressive symptoms in MDD population where the symptoms are present. A potential clinical implication of our findings (i.e. enhanced stress resilience from prophylactic treatment) suggests that riluzole treatment may instead be well-suited for currently remitted MDD or BPD patients who are at high risk for relapse. Several populations may also benefit from potential prophylactic riluzole treatment. These include remitted patients who still display residual symptoms or at high risk of relapse after monoaminergic antidepressant treatment discontinuation [40–43].

Consistent with existing evidence from preclinical and some clinical studies, we demonstrated that riluzole could prevent or block the emergence of specific behaviors related to symptom dimensions of stress-related illnesses, including anhedonia-, and helplessness-like behavior, as well as overall behavioral emotionality. A major limitation of the present work is that it is a male only mouse study. Given that MDD is more prevalent among females [128], and postmortem studies identify more severe cellular deficits in females [129–131], future studies should investigate potential sex-differences in response to riluzole treatment. Although we did not investigate cellular mediators involved in riluzole prophylactic action, this study provides a promising starting point for the use of riluzole for enhancing stress resilience through prophylactic or preventative treatment regimens. Our work raises the possibility of new therapeutic uses for riluzole including as a preventative or prophylactic treatment in individuals experiencing not only chronic stress, but also those subjected to severe acute stressors with high likelihood to develop depressive symptoms.

## Acknowledgements

We acknowledge Discovery Fund fellowship and Ontario Graduate Scholarship for supporting C.F. during the studies. We also acknowledge Labatt Family Network for Research on the Biology of Depression for supporting Y.B. M.B. is supported by the CAMH Discovery Seed Fund and the Canadian Institutes of Health Research (PJT-165852). E.S. was supported by the Brain & Behavior Research Foundation (#25637) and Canadian Institutes of Health Research (PJT-153175). The project was also supported by the Campbell Family Mental Health Research Institute.

## Statement of Ethics

This study protocol was reviewed and approved by [Animal Care Committee and Centre for Addiction and Mental Health], approval number [ACC725]

## Author Contributions

Corey Fee, Yashika Bansal and Keith Misquitta. conducted all experiments, and analyzed data. Yashika Bansal., Corey Fee, Sierra Codeluppi, and Mounira Banasr wrote the manuscript. Mounira Banasr designed and supervised the studies. Etienne Sibille, Robert M. Berman, Vladimir Coric and Gerard Sanacora provided scientific advice on study designs and reviewed the manuscript.

## Data Availability Statement

All data generated/analyzed during this study are included in this article. Further enquiries can be directed to the corresponding author [MB]. A preprint version of this article is available on BioRxiv [132].

